# GABAergic neurons are major contributors of network inhibition in the neonatal hippocampus in-vivo

**DOI:** 10.1101/2025.04.26.650764

**Authors:** Salomé Mortet, Rachel Carayon, Sophie Brustlein, Agnès Baude, Rosa Cossart, Pierre-Pascal Lenck-Santini

**Affiliations:** Aix-Marseille Université, Institut de Neurobiologie de la Méditerranée, INSERM UMR1249 Turing Centre for Living systems, Marseille, France, 163 Avenue de Luminy, Marseille Cedex 09

## Abstract

During development, neural network maturation is activity dependent. During the neonatal period, activity is provided by intermittent, spontaneous network activity patterns (SNAP) that occur independently of environment stimuli. Among the neurotransmitters that take part in neonatal development, the Gamma-amino-butyric acid (GABA) plays a pivotal role. GABAergic cells are the first to emerge the first to form functional synapses. GABA antagonists block hippocampal SNAPs in-vitro and alterations of GABAergic function dramatically impact cortical maturation. Based on these data, the traditional view is that the depolarizing action of GABA, a hallmark of immature networks in slice preparations, is at the core of developmental processes. However, in-vivo evidence for such depolarizing role is not clear, raising questions about the contribution of GABAergic neurons in-vivo. To address this issue we developed an in-vivo approach combining optogenetics and single-unit electrophysiology in non-anesthetized mice. This allowed us to both identify and manipulate hippocampal GABAergic cells while examining their influence on hippocampal SNAP. We found that, even during the first post-natal days, the net action of GABA is inhibitory, but not excitatory. This inhibitory action of GABA drastically increases after the second post-natal week.

## Introduction

The construction of brain networks during early development relies heavily on neuronal activity (Cancedda et al., 2007; Penn et al., 1998; Spitzer, 2006; Wang et al., 2007; Wong Fong Sang et al., 2021). In the neonatal brain, such activity is provided by recurrent, correlated bursts of neuronal firing that emerge independently environmental stimuli and propagate to the cortical areas (Ackman et al., 2012; Babola et al., 2018; Hanganu et al., 2006; Inácio et al., 2016; Karlsson et al., 2006; Leinekugel et al., 2002; Marguet et al., 2015a; Mohns & Blumberg, 2010; Valeeva et al., 2019), where they drive the maturation of targeted networks (Blankenship & Feller, 2010; Cancedda et al., 2007; Hanganu-Opatz, 2010; Katz & Shatz, 1996; Khazipov & Luhmann, 2006; Leprince et al., 2023; Penn et al., 1998; Wang et al., 2007; Wong Fong Sang et al., 2021). Among the neurotransmitters involved in these processes, the gamma aminobutyric acid (GABA) is considered as a key player. Indeed, cortical GABAergic neurons are born and migrate earlier than their glutamatergic counterpart (Dupuy & Houser, 1996; Schlessinger et al., 1978; Tricoire et al., 2011) and GABAergic synapses are the first ones to emerge (Khazipov et al., 2001; Tyzio et al., 1999). Furthermore, alterations of GABAergic function during development affect major aspects of corticogenesis, including cell migration, apoptosis or synapse formation (Bragg-Gonzalo et al., 2024; Cancedda et al., 2007; De Marco García et al., 2011, 2015; Duan et al., 2020; Griguoli & Cherubini, 2017; Manent & Represa, 2007; Mennerick & Zorumski, 2000), they are event considered at the origin of neuro-developmental disorders such as autism spectrum disorders or early epileptic syndromes (Nomura, 2021; Rubenstein & Merzenich, 2003; Tempio et al., 2023; Yu et al., 2006).

Perhaps the most striking argument supporting the role of GABA during development is its direct influence on correlated bursts of activity in neonates. In slices, the immature hippocampus generates its own recurrent bursts, referred to as *giant depolarizing potentials* (Ben-Ari et al., 1989) that involve the collective activation of large numbers of pyramidal neurons (PN) and interneurons (Bonifazi et al., 2009; Khazipov et al., 1997). Importantly, blockers or agonists of GABA_A_ receptors block or promote GDPs, retrospectively (Ben-Ari et al., 1989; Cherubini et al., 1990, 1991; Khalilov et al., 1999; Khazipov et al., 1997) and stimulation of specific GABAergic interneuron subtypes alters GDP frequency (Bocchio et al., 2024; Flossmann et al., 2019; Khalilov et al., 1999; Khazipov et al., 1997; Wester & McBain, 2016). Explaining this facilitating effect on GDPs, an influential theory proposes that, contrarily to adults where GABA is inhibitory, activation of GABA_A_ receptors in neonates induces a depolarization of the neurons. This excitatory action of GABA would originate from the elevated intracellular chloride concentration ([Cl]_i_) observed in neonates, causing an outflow, instead of an entry of chloride ions to the cell (Cherubini et al., 1991; Sato et al., 2017; Sipilä et al., 2005). With age, the progressive increase of extruding chloride transporters (KCC2) would reduce [Cl]_i_, causing a switch of GABA from excitatory to inhibitory.

In-vivo, evidence for the excitatory action of neonatal GABA is not clear. In the immature neocortex, as early as P3, studies showed that although it may be depolarizing, GABA release is not sufficient to excite glutamatergic cells. (Kirmse et al., 2015; Murata & Colonnese, 2020). In the hippocampus, conflicting results are observed. Whole cell recordings of pyramidal cells reveal that the activation of immature GABAergic neurons does not increase but rather inhibit excitatory post-synaptic currents (Valeeva et al., 2016). In contrast, chemogenetic excitation or inhibition of P3 hippocampal GABAergic cells respectively increase or decrease multi-unit activity in-vivo (Murata & Colonnese, 2020). These results being in apparent contradiction, it is still not clear how GABAergic cells influence the activity of immature networks in-vivo.

The immature hippocampus in-vivo is characterized by recurrent patterns of activity called early sharp waves (eSPWs) that reverse in the border of the pyramidal cell layer. Contrarily to GDPs that are generated locally, eSPWs have an external origin: they are triggered by myoclonic movements that activate somatosensory and entorhinal cortices and propagate to the hippocampus (Dard et al., 2022; Graf et al., 2021; Karlsson et al., 2006; Khazipov et al., 2004; Marguet et al., 2015a; Mohns & Blumberg, 2008; Sokoloff et al., 2021; Valeeva et al., 2019). While there is evidence for GABAergic activity during eSPWs (Dard et al., 2022; Leinekugel et al., 2002) the precise timing of this activity is not known.

To better understand how GABAergic neurons contribute to eSPWs and influence hippocampal networks in-vivo, we recorded single unit activity in unanesthetized neonatal mice during the first two post-natal weeks. Adapting the photostimulation-assisted identification of neuronal population (PINP) method(Lima et al., 2009) to neonatal mice, we were able to identify and manipulate the activity of GABAergic neurons. In general, GABAergic neurons were more active than the rest of the recorded population and participated to more eSPWs. We found evidence of both spontaneous and GABA-driven inhibition as early as P3. This inhibitory effect increased rapidly at the beginning of the 2^nd^ post-natal week to reach a plateau at P9-P10. Altogether, our results suggest GABA is principally inhibitory the neonatal hippocampus in-vivo and that this inhibition is not mature until the 2^nd^ post-natal week.

## Results

Local field potentials (LFPS) were recorded bilaterally from the dorsal CA1 region of the hippocampus in P3 to P12 head restrained mouse pups (Figure 1A). As reported previously (Karlsson & Blumberg, 2003; Leinekugel et al., 2002; Marguet et al., 2015b; Valeeva et al., 2019), active sleep in neonates was characterized by an isoelectric hippocampal LFP interrupted by eSPWs, occurring at a rate of 2.97 ± 0.16 events/min (Figure 1B, C). ESPWs were often followed by a 0.5-2s period of delta (2-4Hz)-beta (10-20Hz) oscillations coupled with increased action potential activity (the ‘tail’ period Figure 1C).

**Figure 1.**
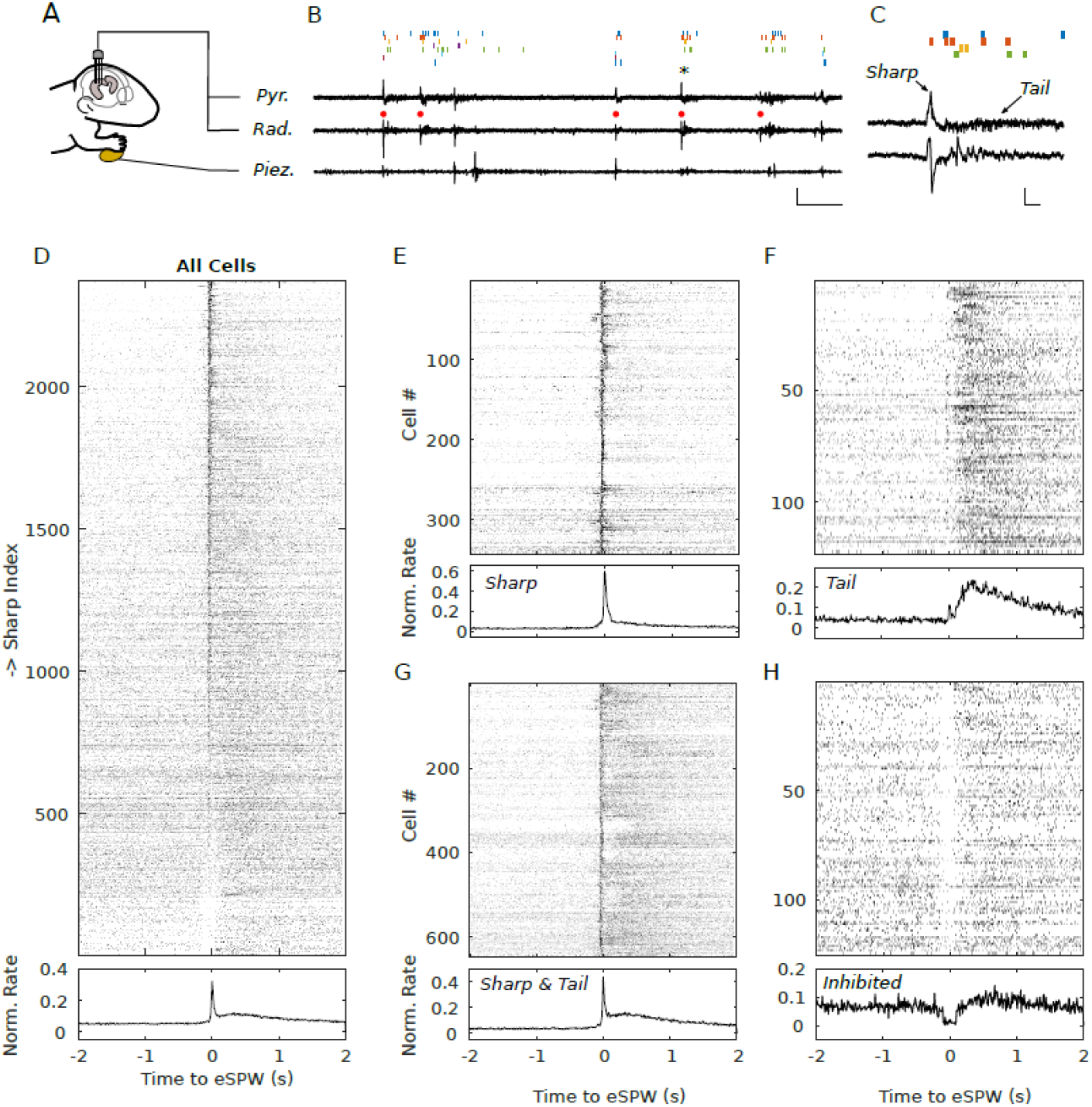
Electrophysiological recordings in head-fixed, non-anesthetized neonatal mice. A) Illustration of the recording principle. B) Example of LFP traces recorded in the pyramidal cell layer (Pyr.) and *stratum radiatum* (Rad), together with the movement outputs from one piezo-element glued to a forepaw. Vertical bars of different colors indicate the occurrence of APs from different cells, isolated from the spike sorting process. Red dots indicate eSPWs (calibration bars: vertical: 200 µV; horizontal: 5s). C) Close-up view of the traces at the time surrounding the eSPW indicated by an asterisk (calibration bars: vertical: 100 µV; horizontal: 200ms). D) Top: Cell response to eSPW from the entire dataset. Each line represents the PETH of an individual cell, normalized by the peak firing rate in the PETH and expressed as a linear grayscale (white=no firing, black=maximum firing). Cells are sorted by decreasing SI. Bottom: average for all cells. E-H) Similar representation selecting cells from four classes of responses.

### Single unit activity in CA1

Using template matching spike sorting algorithms (Yger et al., 2018), we isolated the firing activity of 2380 neurons in a total of 293 sessions recorded from 98 pups. Each pup was recorded on a single day. Pooling recordings from P3 to P12, we first investigated the properties of action potential (AP) waveforms (Supplementary Figure 1), extracted from LFP recordings (100-5000Hz). Waveform duration, measured at 25% of the maximum spike amplitude, was large, with an average of 675.0±3.0 µs. By comparison, pyramidal cells recorded in 6 supplementary juvenile, P27-P30 mice, under the same conditions (N=396 cells) reached a significantly shorted duration (553.9±6.9 µs; t=17.47, p=3.95*10^-65^). Also, the firing rate of neonatal hippocampal neurons was extremely low (mean: 0.50±0.016 AP/sec) with prolonged periods of silence interrupted by eSPWs. In comparison, P27-30 mice fired at significantly higher rates (mean=2.32±0.19 AP/sec; t=-22.62, p=1.15x10^-104^). In contrast with adult neurons and data from our P27-30 mice, it was not possible to identify GABAergic neurons on the sole basis of spiking properties. Indeed, scatter plots displaying spike width, firing rate and pre/post AP hyperpolarization did not reveal any identifiable cluster. In contrast, such cluster was identifiable in juveniles (Supplementary Figure 1D and E). In the following section, we will therefore consider all units as part of a single group.

### Influence of eSPWs on single unit activity

To better quantify the influence of eSPWs on single unit firing, we performed rasters and peri-event time histograms (PETHs Supplementary Figure 2) for each cell, considering the time of the eSPW trough as the reference event. Data from P3 to P12 animals were pooled. Confirming previous reports in rats (Mohns & Blumberg, 2010; Valeeva et al., 2019), we observed that a large proportion of mouse CA1 neurons increased their firing rate at the time of eSPWs. The type of response displayed by the neurons was heterogeneous: some fired specifically at the trough of the eSPW some at the tail and other during both times. While most of these neurons did not fire in every sharp (Supplementary figure 2 ), the response pattern they exhibited when they were active was always the same. To better visualize single unit activity at the population level, we grouped the PETHs of all recorded cells on Figure 1D. It can be noted that there is a global, sharp increase of firing at the time of the eSPWs, with a peak in the 0ms bin of the averaged PSTH. The distribution of peak firing times is at its maximum in the 0ms bin (Supplementary Figure 3, median at 31.5ms) and the mean Sharp Index (SI, see methods) significantly differs from 1 (m=3.83, t-test on log-transformed data, excluding 199 cells with SI=0; t_2181_=39.89; p<0.0001; Supplementary Figure 3), meaning that neurons fire more during the *sharp* period than during the rest of the session.

We then estimated whether cells systematically fired during the sharp (from 0.1 s before to 0.1 after the eSPW) and/or during the tail (0.2 to 1.5 s after the trough -paired t-tests p<0.05) components of eSPWs. This allowed us to identify several classes of neurons (Figure 1E-H): Neurons that were not systematically modulated by eSPWs (n=1267, 53.4% Supplementary Figure 4), neurons that significantly increased their firing rate during the sharp period (14.5%, n=343; Figure 1E); during tails (5.2%, n=123; Figure 1F), during both sharp and tails (27.1%, n=643; Figure 1G). Note that, even for cells that did not reach significance, a positive modulation can also be observed in the 0-10 ms time bin. Importantly, we observed a fifth class of cells that systematically decreased their firing rate during the sharp component (5.3%, n=125; paired t-test, p<0.05; Figure 1 H). This class was not exclusive since some of those cells also belonged to the *tail* class: they were inhibited during the sharp but increased their firing during the tail component. The presence of inhibition during eSPWs is also quantifiable with the SI (Supplementary Figure 3), where values below 1 correspond to a decreased activity (n=616), and values at zero correspond to no firing at all (excess numbers in the zero-bin, n=199). These cells are also visible at the bottom of Figure 1D, where they form a white, vertical band surrounding the eSPW. It is therefore likely that these neurons were inhibited during eSPWs.

### Effect of age on eSPW single unit response

We then examined whether the cell properties evolved with age. As pups grew older, the impact of eSPWs on neuronal firing decreased (Figure 2). In parallel, the session firing rate increased significantly with age (Age vs. log(Session-Rate) correlation coefficient: r=0.480, p=1.04*10^-137^, confirmed by Generalized linear mixed-effects models -GLMM-including animal ID as a variable Supplementary data; Figure 2B) and SI, which also depends on total firing rate decreased with age (Age vs. Log(SI); excluding inhibited cells; r=-0.463, p=1.21*10^-115^, also confirmed by GLMMs; Figure 2 C). As can be observed in Figure 3 A, this is likely caused by both an increase of the background activity and a decrease in eSPWs firing with age. By P9, the ESPW-associated PETH barely reached values above baseline fluctuations. This phenomenon is also illustrated (Figure 2D) by a progressive decrease of the proportion of cells that were significantly activated during eSPWs (determined using paired t-tests). Importantly, the proportion of neurons inhibited during eSPWs increased with age (Figure 2E). Altogether, these data suggest that inhibition is present during the first post-natal week and that it increases with age. The apparent antagonistic age-related behavior between eSPW excitation and inhibition suggests that it is the emergence of GABAergic inhibition that causes the decrease of eSPW-associated firing.

**Figure 2.**
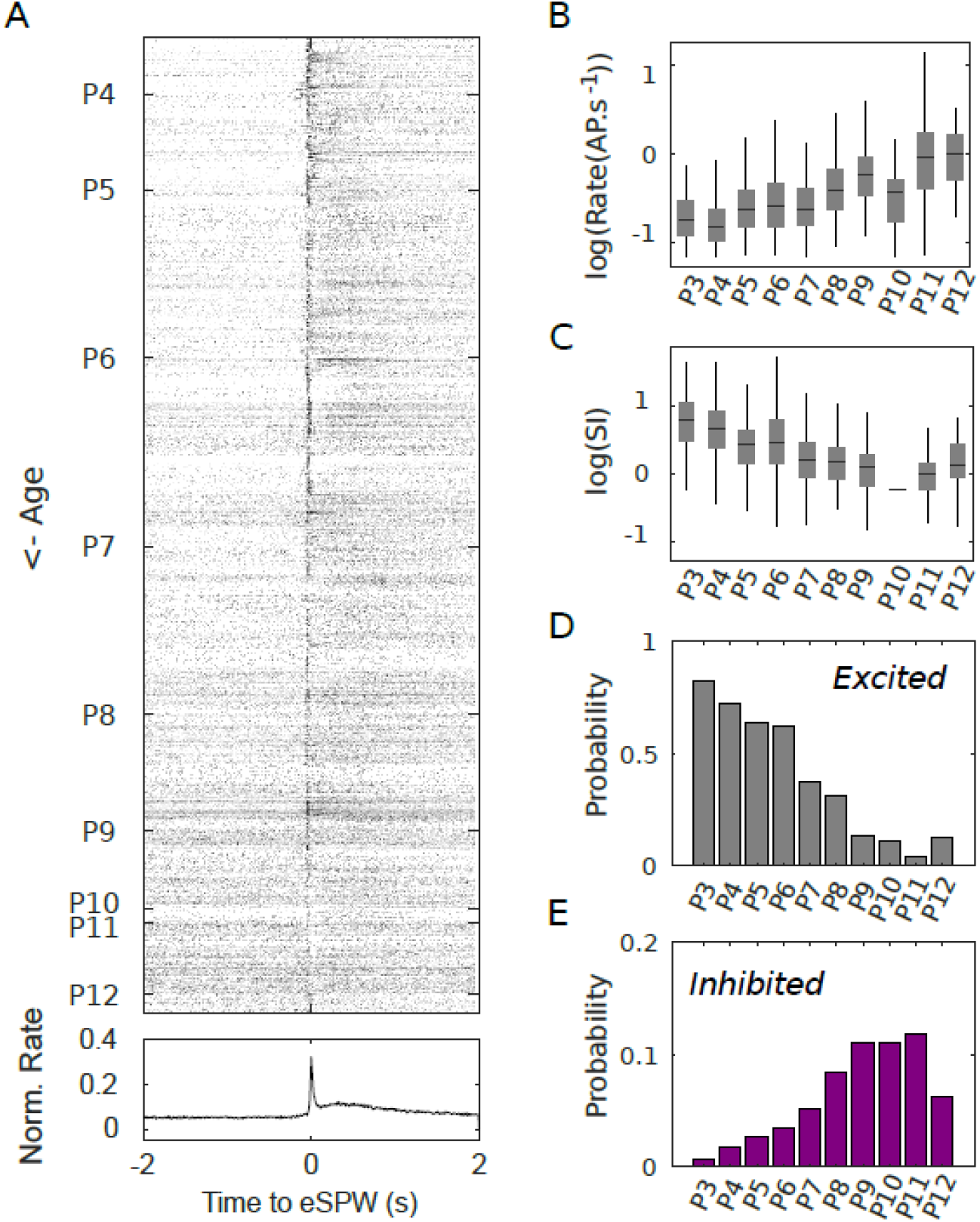
Evolution of firing properties as a function of age. A) PETHS for the whole dataset with cells sorted by increasing age. B-C) Box plots of log-transformed of firing rates and SI as a function of age. Horizontal bars: median, lower and higher extremities: 25^th^ and 75^th^ percentiles, vertical bars: limit for outliers. D-E) Evolution of the proportion of cells that are excited or inhibited during eSPWs, respectively.

**Figure 3:**
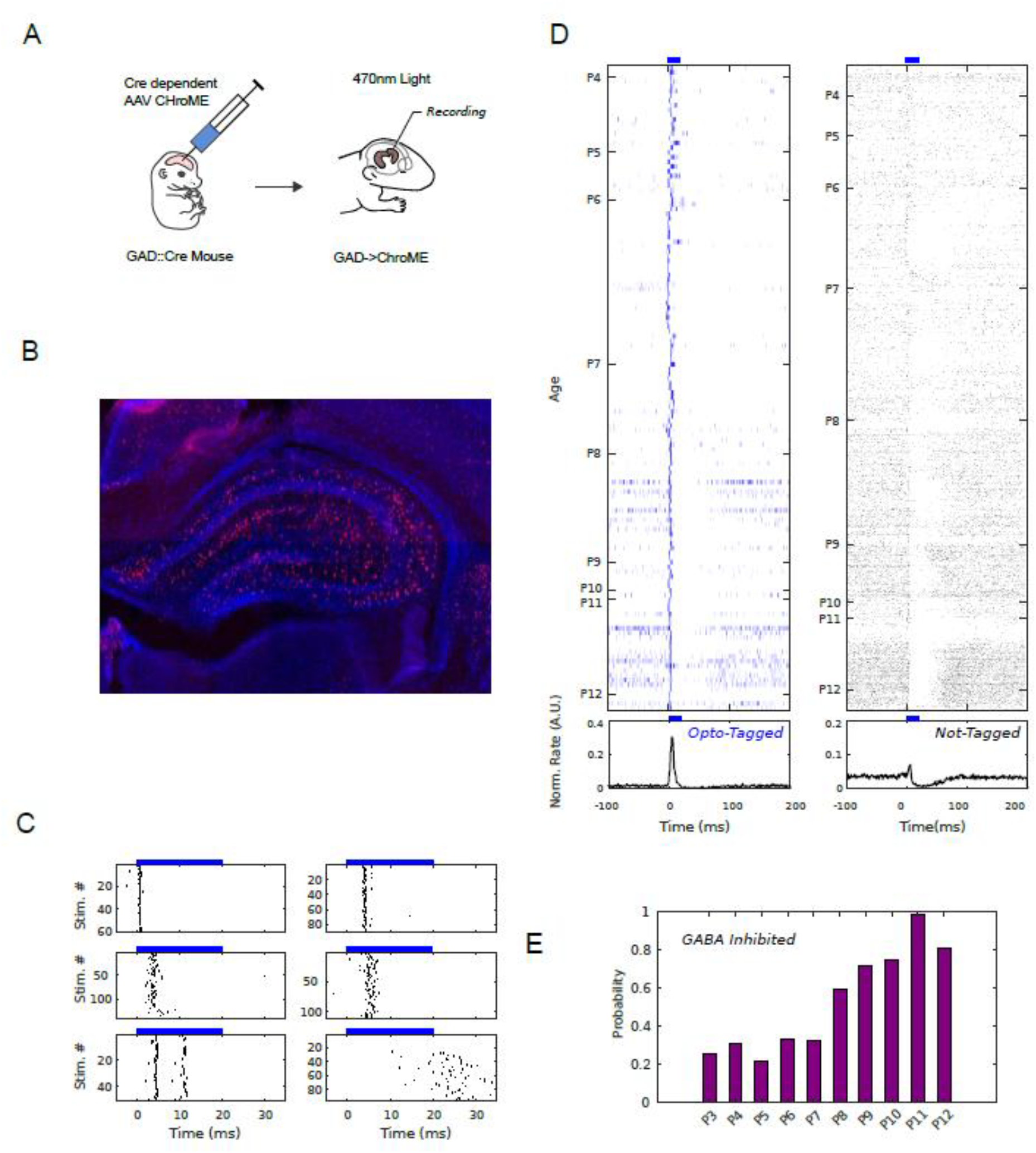
Opto-tagging of GABAergic cells and indirect network inhibition. A) GAD-Cre mice received ICV injections of ChroME expressing AAVs either at E16 or at P0 for recordings at P3-P6 or later, respectively. B) Coronal hippocampal section of a P3 mouse demonstrating de presence of reporter fluorescent protein mRuby (red) and DAPI (blue). C) Rasterplots of 6 different cells with fast delay (top four plots), rebound firing (bottom left) and late onset (bottom right). Dark dots represent action potentials of the cell, each row a stimulation trial. Blue rectangle: timing and duration of the photo-stmulation. D) Average PSTHS of tagged and not tagged cells sorted as a function of age. Bottom: population average. Note the decrease of firing during and after photo-stimulation. E) Proportion of cells that are significantly inhibited after photo-stimulation of ChroME-expressing GABAergic cells.

### Optogenetic identification and manipulation of immature GABAergic cells

Since it is not possible to dissociate pyramidal cell from interneuron AP waveforms, we therefore adapted an optogenetic method developed for adults (Lima et al., 2009) that could both help detect and manipulate GABAergic cells. Briefly, adeno-associated viral vectors (AAVs) containing sequences (*pAAV-CAG-DIO-ChroME-ST-P2A-H2B-mRuby3*) expressing the opsin ChroME were injected intraventricularly in Gad::Cre expressing pups. To allow sufficient protein expression before recordings, injections were performed in 51 pups, either in-utero (at Embryonic stage E15), for P2 to P5 recordings or at P0 for recordings that started at P6. At the end of recordings, we injected a series of short (20 ms), blue light (270 nm) pulses with either a LED or a laser (for higher power stimulations, see methods), via a fiber that was attached to the electrode array shanks.

Of a total of 1492 recorded cells (also belonging to the previous dataset but restricted to session where photo-stimulation induced a visible effect on the LFP), 131 (8.8%) were directly (within 10ms) and repeatedly (more than 5 stimulations) excited by the light stimulation (Figure 3). These cells will be referred to as ‘*opto-tagged*’ cells. Among these cells, approximately half displayed a narrow, 1-2ms excitation window Figure 3C). Nine of these cells fired twice, in rebound, with a systematic interval ranging between 10 and 20 ms depending on the cell (Figure 3C -bottom left). Other opto-tagged neurons responded at more variable times, displaying a wider excitation window ranging from 5 to 20 ms (Figure 3C-middle row). Interestingly, 6 neurons (recorded between P4-P7) fired more than 10ms after the light onset (Figure 3C bottom right). It is unfortunately impossible to determine whether these cells were directly depolarized by the light (when they fired within the 20ms light stimulation), whether they were indirectly excited by other neurons or fired by disinhibition after GABAergic cells stopped firing.

Using histological reconstruction of the recording sites, the presence of APs on specific electrode contacts and the distance separating these contacts to the site of inversion of eSPW LFPs (in the superficial part of the str. pyramidale), it was possible to estimate the localization of opto-tagged cells with regards to the pyramidal cell layer. As seen on Supplementary Figure 6, opto-tagged cells were likely recorded in all recorded CA1 layers, including the str. oriens, the str. pyramidale, the str rad. and str lac. mol.

### Stimulating opto-tagged cells induces inhibition in neonates

The PSTHs of neurons that did not systematically fire during light stimulation are displayed in Figure 3D. From this graph, two major observations can be made. First, there is an excess firing that peaks 5 ms after stimulus onset. This short delay suggests that this increase is caused by direct photo-stimulation, likely involving GABAergic cells that do not express ChroME in large enough quantities to systematically fire after light stimulation. Indeed, we identified 54 cells that fire in a low number of trials (less than 5) but display an increased rate at stimulus onset. Second, following this 5 ms peak and lasting for ∼40ms, there was a systematic decrease of the activity of a large number of cells. Statistical comparisons between baseline and post-stimulation on a trial-to-trial basis revealed that this inhibition concerned 42.8% of cells (638 neurons with significant paired t-tests). This inhibition was observed both in opto-tagged (n=65 over 131, 26.7%) and non-tagged (n=573 over 1361, 42.1%) neurons (proportions not significantly different, X^2^= 2.7587. p=0.096). To better estimate the developmental effect of GABAergic stimulation on the network, we restricted our dataset to laser stimulations, which provide a stronger power and illuminated cells from all shanks at a time (35 pups, 1492 cells). Inhibition following photo-stimulation was observed as early as P3, where it affected 25.6% of cells (Figure 3E). This proportion increased sharply at P8 and reached 100% at P11 (Figure 3E).

### Significant recruitment of GABAergic neurons during eSPWs

As seen on Figure 4A opto-tagged cells displayed similar response profiles than the rest of the population recorded in the same conditions (Figure 4B, Not-tagged cells). Opto-tagged cells also increased their firing rate at the trough of the eSPW and they also displayed a variety of responses similar to those observed in non-tagged cells. That said, their firing rate (mean=1.41±0.186Hz) was higher than for non-tagged cells (Figure 4C; mean=0.52±0.186Hz, ttest on logtransform data, t_1490_=6.64; p=4.02*10^-14^) and they fired in a larger proportion of eSPWs than non-tagged cells (Figure 4D; Opto-tagged: mean= 43.6.±2.84%, Not-tagged: mean=31.29±0.67%; t-test on the arcsin transform of the proportions: t=5.24, p=1.82*10^-7^). Considering *Sharp* and *Sharp+Tail* cells only, the average spike time during the sharp period (±100 ms around the eSPW trough -Figure 4E) was smaller for opto-tagged neurons (mean=9.9±2.86ms) than for not-tagged neurons (mean=16.3±0.96ms, t=2.098, p=0.037). Furthermore, the activity of opto-tagged cells was more likely to be significantly modulated by eSPWs than not-tagged cell (49.6% vs 39.6% of modulated cells in opto-tagged and non-tagged cells, respectively; χ^2^ test of independence χ ^2^_(1, N=1492)_=4.98, p=0.026). Importantly, 10.7% of opto-tagged cells (n=14) were significantly inhibited during eSPWs, suggesting that they also received inhibitory inputs. SIs were similar between the two classes of cells (Figure 4F). Altogether, these results suggest that GABAergic neurons are strongly active during eSPWs, and therefore contribute significantly to the inhibitory activity observed during eSPWs.

**Figure 4.**
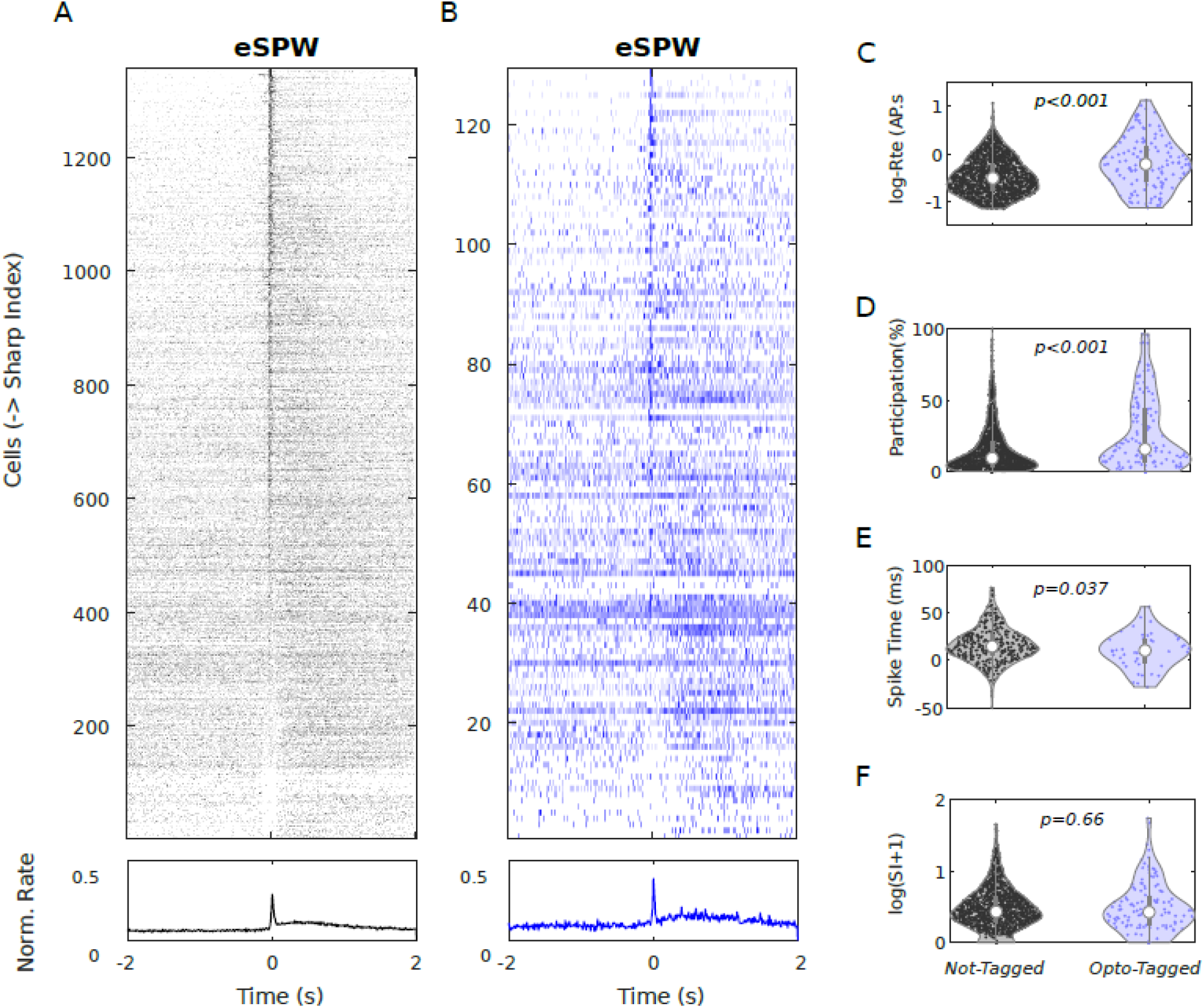
Firing properties of opto-tagged neurons. A) PETHs of not-tagged and B) opto-tagged neurons together with the population average (bottom). C-F) Violin plots of firing measures for both types of cells (kernel density estimates). Each dot is the measure for an individual cell. White circle: median; grey, thick vertical bar: 25^th^ and 75^th^ percentile, thin vertical bar: outlier limit. p: p-value.

## Discussion

Using a novel method based on opto-tagging of hippocampal GABA neurons in vivo, we were able to show that these cells were highly active during the first post-natal week and that they exert an inhibitory action on approximately a 3^rd^ of the recorded neurons. Evidence of putative GABA-drive excitation at this age was inconsistent and only concerned a negligible (∼0.4%) of recorded cells. In accordance with these results, single unit recordings also showed evidence for eSPW-driven inhibition as early as P3. This proportion increased with age, so did the proportion of cells inhibited after photo-stimulation of ChroME-expressing GABAergic neurons. After a rapid increase at the beginning of the 2^nd^ post-natal week, it reached 80-100% at P10-P12. Altogether, these results suggest that endogenous GABA is inhibitory in the neonatal hippocampus and that its inhibitory influence sharply increases during the 2^nd^ Post-Natal week.

### Relevance of the method

Since the discovery of GDPs and the shift in intracellular chloride concentrations during development, a major challenge has been to characterize the role of GABAergic activity in immature networks in-vivo. By adapting traditional in-vitro techniques to live, immature animals, investigators have been able to reveal, in pyramidal cells, the simultaneous presence of glutamatergic and GABAergic (GABA_A_) currents during eSPWs (Leinekugel et al., 2002), reminiscent of what happens during GDPS. However, such techniques only provide data from a small number of cells at a time, preventing experimenters to grasp the full diversity of neuronal activity according to cell type or age. To this day, there is no report of intracellular activity in immature GABAergic neurons during eSPWs in-vivo.

Recently, exciting results have been produced by calcium imaging recordings in live immature mice. Kirmse et al. (2015), showed that, although exogenous GABA delivery depolarized cortical plate neurons at P3-P4 in-vivo, it failed to induce calcium transients. The authors concluded that GABA was depolarizing but not excitatory in the immature neocortex in-vivo. More recently, Dard et al. (2022) investigated immature calcium transients in both hippocampal pyramidal and GABAergic neurons in-vivo. They showed that during the 1^st^ post-natal week, hippocampal pyramidal cells and interneurons were synchronously activated following myoclonic twitches and startles. At the beginning of the 2^nd^ post-natal week, pyramidal cells, but not interneurons disengaged from movement. This discontinuity in hippocampal dynamics, coinciding with the emergence of peri-somatic inhibition, suggests that the inhibitory action of GABA only impacts hippocampal networks after the 2^nd^ post-natal week. However, the temporal dynamics of calcium fluorescent indicators are significantly slower than electrophysiological events such as APs. This may be particularly problematic for fast inhibitory activity, which may not be detected when surrounded such an intense AP firing as the one observed during myoclonic twitches and eSPWs. Despite remarkable advances in calcium indicators and spike inference algorithms (Berens et al., 2018; Deneux et al., 2016; Vogelstein et al., 2010) there are still functional differences between neuronal activity recorded in electrophysiology vs calcium imaging (Siegle et al., 2021; Wei et al., 2020). Therefore, current calcium fluorescence techniques may not reveal the full activity repertoire of immature GABAergic neurons in-vivo, particularly during the first post-natal week. As we saw in our extracellular dataset, it is almost impossible to discriminate between interneurons and pyramidal cells on the sole basis of extracellular waveform and/or discharge properties. For this reason, we developed the opto-tagging method to identify GABAergic neurons within our single unit recordings. This allowed us to investigate neuronal activity at the millisecond resolution while keeping track of the identity of GABAergic neurons. Notably, we observed an heterogeneity of responses during eSPWs, including a short-lasting (∼100ms) inhibition in a small portion of cells. This was not observed in calcium imaging data.

### GABAergic Inhibition but not excitation in the immature hippocampus

Our results provide evidence for an inhibitory role of GABAergic neurons during the first post-natal week in-vivo. In addition, less than 1% of recorded neurons fired with a long delay (15-25 ms) after the photo-stimulation onset. While these neurons (all recorded before P8) could have been excited by GABAergic transmission, their response delays were rather short compared to what is observed in-vitro (∼40ms), under physiological, immature intracellular chloride concentrations (Valeeva et al., 2010). In any case, the low proportion of these cells (∼0.4%) suggests that, even during the first post-natal week, the excitatory action of GABA on hippocampal single unit firing is negligeable compared to its massive inhibitory effect on the network. Therefore, although acute slice preparations experiments support the excitatory GABA hypothesis, our result fail to do so in-vivo. Yet, recent evidence suggests that immature GABA may be indeed depolarizing in-vivo. First, [Cl]_i_ is elevated in immature neurons in-vivo (Sato et al., 2017). Second, exogenous GABA induces a depolarization of the majority of cortical plate neurons at P3-P4 (Kirmse et al., 2015). Finally, using chemogenetics Murata and Colonnese (Murata & Colonnese, 2020) showed that excitation or inhibition of hippocampal GABAergic cells increased or decreased neuronal firing in vivo. To reconcile these results with ours, one may consider the depolarizing action of GABA as facilitating, rather than excitatory. Indeed, the fact that, in neonates, the reversal potential of GABA (∼-60µV) sits close to the AP threshold of the cell (∼-45 µV) makes it a good candidate to facilitate ongoing glutamatergic inputs and indirectly trigger APs. This scenario likely occurred in Murata et al. (2020) experiments where spontaneous multi-unit activity was studied in the context of prolonged activation of GABAergic cells. In contrast, our photo-stimulations triggered short bursts of GABA release, occurring randomly during our recordings. Since background activity in neonates is really low, GABA release was more likely to be triggered alone, without the co-occurrence of glutamatergic inputs. As demonstrated by Kirmse et al., (2015) and explaining the absence of GABA driven excitation in our data, GABA release by itself is not sufficient to induce AP firing. While our experimental procedures do not allow us to investigate subthreshold events, recent advances on voltage sensitive dyes may help us reveal such effects.

### Origin of early Gabaergic inhibition

During the 1^st^ post-natal week, we observed a consistent suppression of AP firing in a third of the hippocampal units. This inhibitory effect may be caused by a GABA-induced hyperpolarization or by a shunting effect of GABA, mechanism through which glutamatergic inputs are cancelled by both a reduction of input resistance of the cells and a re-entry of chloride ions when its membrane potential reaches the GABA reversing potential (Gao et al., 1998). Another, non-exclusive possibility is that the co-activation of metabotropic GABA_B_ receptors induced a significant inhibitory effect. This effect has indeed been demonstrated in slices (Khalilov et al., 2017; Tosetti et al., 2004, 2005).

Given the late rise of perisomatic inhibition from PV basket cells (Dard et al., 2022), such neonatal inhibition may originate from dendrite targeting GABAergic interneurons. While we cannot determine precisely the type of GABAergic cell we recorded, our LFP/histological reconstructions suggest that opto-tagged cells likely belong to interneurons with cell bodies in multiple CA1 layers, including but not restricted to the pyramidal cell layer. All cell types are therefore likely to contribute to inhibition at these ages. Further studies may reveal the differential contribution of specific subclasses GABAergic cells and receptors to early inhibition.

### The rise of inhibition during the second post-natal week

The proportion of cells inhibited by photo-stimulation of GAD-Cre neurons rose-up sharply, going from ∼20% between P3 and P7 to 60% at P8 and even 100% at P10. Similarly the proportion of cells inhibited by eSPWs increased. In the same time, the excitatory impact of eSPWs on the network decreased. These results are in agreement with a recent study from Dard et al. (2022) that showed a rapid rise of the movement-induced inhibition during the second post-natal week. This rise of inhibition was paralleled with the emergence of peri-somatic inhibition by Parvalbumin-expressing GABAergic cells. In addition to the maturation of peri-somatic inhibition, other developmental processes, such as the maturation of GABA_B_ signalling (Luhmann & Prince, 1991) or the upregulation of the chloride extruder KCC2 (Rivera et al., 1999) may participate to such an increase.

## Conclusion

Altogether, our results provide strong evidence for a net inhibitory effect of GABA signalling in the neonatal hippocampus. Is also confirms previous reports of a rapid rise of inhibition after the second post-natal week. As argued in Dard et al. (2022) and (Kirmse & Zhang, 2022), it is likely that the development GABAergic inhibition at the beginning of the 2^nd^ post-natal week allows CA1 neurons to disengage from sensory inputs driven by the periphery such as myoclonic twitches/startles and retinal or cochlear waves. This late disengagement from the periphery may allow the emergence of internal hippocampal dynamics, necessary for the initiation of environment exploration.

## Methods

All experimental procedures, husbandry and care were approved by the French ethics committee (Ministère de l’Enseignement Supérieur, de la Recherche et de l’Innovation (MESRI); Comité d’éthique CEEA-014; APAFiS #28.506 #46290-2023121410536843 v5 and S #38060-2022072115429698 v9). They were conducted in agreement with the European Union Directive 2010/63/EU on the protection of animals used for scientific purposes.

Mice were bred and stored in an animal facility with room temperature (RT) and relative humidity maintained at 22 ± 1°C and 50 ± 20%, respectively. They were provided ad libitum access to water and food and housed in enriched social and environmental conditions.

GAD67-Cre mice were kindly donated by Dr. Hannah Monyer (Heidelberg University) and back-crossed to a JAX Swiss Outbred background. GAD-Cre males were then bred with wildtype Swiss females.

Viral vectors (pAAV-CAG-DIO-ChroME-ST-P2A-H2B-mRuby3), obtained from Addgene and donated by H. Adesnik (viral prep # 108912-AAV9; http://n2t.net/addgene:108912; RRID:Addgene_108912) were used to express the cre-dependent cation channelrhodopsin ChroME and target it to the soma and proximal dendrites of neurons (Mardinly et al., 2018). The opsin was also separated from the reporter protein mRuby3, which remained in the nucleus. During the initial stages of the experiment, two litters (7 animals P5-P7) were injected with another viral vector (pAAV-EF1a-double floxed-hChR2(H134R)-mCherry-WPRE-HGHpA (AAV9)) expressing the humanized channelrhodopsin H134R mutant fused with mCherry. In our hands, the light activation induced using this viral last vector was less efficient than with the ChroME-mRuby3 vector.

Depending of the desired recording age (before or after P6) viral vectors were either injected in-utero at embryonic age 15 (E15) or intra-cerebro-ventricularly at P0, respectively.

The intra-cerebro-ventricular injection protocol was adapted from published methods (Bocchio et al., 2024; Kim et al., 2014; Leprince et al., 2023). P0 Gad67cre/+ mouse pups, genotyped the same day, were anesthetized by hypothermia. They were placed on ice and maintained on a dry ice-cooled stereotaxic adaptor (Stoelting, 51615) fixed to a stereotaxic frame with a digital display console (Kopf, Model 940). Solutions (2µL) containing diluted AAV with Fast Blue dye (1:20) were injected in the left lateral ventricle, estimated to be at two fifth of the imaginary line between the left eye and lambda. Injections were performed with a glass pipette pulled from borosilicate glass (3.5” 3-000-203-G/X, Drummond Scientific) and connected to a Nanoject III system (Drummond Scientific). The pipette was inserted 0,4 mm below the surface of the skull. Injections accuracy was visually verified by transparency when the blue dye spread through the whole ventricle. After injection, pups were placed with their litter and recording procedures started 5 days later.

In utero injections (adapted from (Cloarec et al., 2016; Walantus et al., 2007)) were performed on pregnant dams at E15,5 under analgesia (buprenorphine-Buprecare® -0,045mg/kg and carprophen, Rimadyl® -10mg/kg) and general anesthesia (Isoflurane 1,5-2%). The uterine horns were exposed and a volume of 1,5–2 µL of viral solutions with the same concentrations was injected into the left lateral ventricle of each embryo. Injections were performed through a pulled glass capillary with a microinjector (Picospritzer II, General Valve Corporation, Fairfield, NJ, USA). Mice give birth 4 days after the procedure. At P3, the expression of mRuby3 in Gad67cre/+ pups was detectable when mice were placed under green fluorescent light. This allowed us to restrict our recording procedures to Gad-Cre/ChroMe-mRuby3 pups and therefore reduce the number of animals used for experiments.

Validation of the GAD67-Cre transgenic mouse line and ChroME viral vectors, previously done in our laboratory and published in a recent article (Bocchio et al., 2024). Briefly, immunolabelling of injected pups showed that the distribution of mRuby3 expressing cells was consistent to the known distribution of GABAergic neurons in CA1. We did not find any neuron with pyramidal-like structure (with dendrites reaching to the *str. Lac. Mol*) expressing mRuby3. Finally, all mRuby3 expressing cells were GAD67 positive.

Surgery was performed in mice aged between P3 and P12 under isoflurane anesthesia (2-4 %). Before surgery, mice received a subcutaneous injection of buprenorphine (Buprecare® 0,03mg/kg) and topical administration of lidocaine/prilocaine ointment (5 %) on the scalp. During anesthesia, the scalp was scrubbed with pividone-iodine solution(Betadine® 7.5%) and incised with a scalpel. The skull was cleared of periosteum and blood. Two chlorided silver wires (5cm long, 127µm diameter, AM-System), serving as ground and reference, were placed for 1mm between the skull and the dura above the posterior part of the auditory cortex and secured with cyanoacrylate glue (vetbond, 3M). A 3 by 1 cm head bar was then fixed above the cerebellum and posterior visual cortex with dental cement (Superbond, Sun Medical CO.), letting the areas above both hippocampi accessible. Craniotomies were then performed above the hippocampi in the coordinates described on table 1 s a function of age.

**Table 1.** Stereotaxic coordinates for hippocampal electrode insertion (with regards to Bregma) as a function of age.

|  | P3-P4 | P5-P6 | P7-P8 | P9-P10 | P11-P12 |
| --- | --- | --- | --- | --- | --- |
| AP | 1.1mm | 1.3mm | 1.5mm | 1.7mm | 1.9mm |
| ML | 1.3mm | 1.5mm | 1.7mm | 1.9mm | 2.1mm |

The craniotomy sites were covered with agarose (Sigma-Aldrich, 2% in PBS solution) for protection until recording time. Pups were then let to recover without anesthesia on a heating pad. Thirty minutes after surgery, pups were secured to the recording apparatus via the head bar, the rest of the body resting on a heating pad to maintain body temperature at 37°C. Throughout the recording that follows, pups were constantly monitored and fed with veterinary milk ad libitum. A piezo element (RS Pro, 35mm Ø) functioning as a movement detector was placed bellow the animal and connected to custom made electronic amplifier and rectifier montage (https://mptek.fr/).

Electrodes consisted of silicon probes that were previously coated with a fluorescent marker (DiI-Sigma-Aldrich) and secured to micro-manipulators (IVM 300; Scientifica) by specific adapters (Adpt-A32-OM32 and Adpt-A64-OM32x2-sm, Neuronexus) and driven through a computer interface (LinLab2, Scientifica). Two types of silicon probes were used simultaneously. First, 32-channel probes (A1x32-Edge, Neuronexus) with contacts spread linearly along a single axis (50µm spacing covering 1550 µm) were used to detect eSPW LFPs. They were inserted throughout CA1-Dentate gyrus axis of the left dorsal hippocampus. Second, high density 32- or 64ch probes (Buzsaki32, Neuronexus or ASSY-77 P-1, Cambridge Neurotech), mounted with optic fibers were used for electrophysiology and optogenetics. These probes, also coated with DiI were slowly inserted through the right dorsal hippocampus by steps ranging from 20 µm, for the first 800 µm to 1 µm when approaching the pyramidal cell layer where APs appeared. Once units of sufficient amplitude (3 times the background activity) were detected, the electrode was left in place for 15 minutes and recordings started. Once a recording was achieved, the single unit electrode array was moved ∼50-100 µm in order to detect other units below the pyramidal cell layer, identified by the LFP inversion during eSPWs. The signal from the electrodes was filtered (0.1 Hz-6000 Hz), digitized (30KHz) and multiplexed by preamplifiers (RHD-32 channel, Intan Technologies). It was then recorded via an acquisition system (RHD2000_evaluation_system, Intan Technologies) that synchronously recorded the analog signals from the piezo sensors and the TTL ouputs from the optogenetic controlling devices. Signal acquisition was controlled via the Open-Ephys graphical user interface (https://open-ephys.org/gui; Versions 0.6.0 to 0.6.6).

Optical stimulation was performed via optic fibers located either on each shank of the probe (Buzsaki32, neuronexus, 105 μm core fiber, 0.22 NA) or one fiber above all shanks (Assy-77, Cambridge Neurotech -100 µm core fiber, 0.22 NA with 0.7mm taper tip). A critical part of the stimulation protocol (Royer et al., 2010, 2012) is the ability to modulate the intensity of the light pulse so that it progressively increases and decreases (such as between the troughs of a sine wave). To do so we first used four LEDs (470nm, 170mW; Thorlabs) collimated through a lens to optic fibers (100 µm core, 0.22NA) themselves connected to the electrode array with a ferrule. At the extremity each shank fiber, light power reached a maximum of 40µW, which is sufficient to induce AP firing near electrode contacts, but not to excite neurons outside of this region. However, after multiple use of the electrode arrays, we realized that despite careful cleaning, fibers become opaque, and light output becomes dimmer. Under these conditions, it was therefore difficult to induce APs. In later parts of this experiment, we therefore used a more powerful laser source (Sapphire 488, Coherent), that we controlled to output between 110 and 900 µW.

Light sources were modulated using a custom made MATLAB graphical user interface from which we selected the shank excited (In the case of Neuronexus probes) and the light parameters being used. The output from this interface was sent to a Digital/Analog I/O interface (National Instruments USB-6341) and dispatched between LEDs or the laser and the Open-Ephys acquisition board. Using a photo-sensor connected to an oscilloscope, we determined the delays between the emission of the TTLs and the appearance of light on the tip of the electrode. These delays were different between the LED (2ms) and the laser (1ms), they were taken in consideration when computing the cellular responses to photo-stimulation.

Spike sorting was performed offline using the Spyking-circus toolbox, developed in python (Yger et al., 2018). This toolbox uses template a matching algorithm, that allows to retrieve waveforms even in the case of temporal collision with other waveforms from other neurons. This is particularly relevant in periods of high firing such as eSPWs or optogenetic stimulations. Curation and visualization of spike sorting output was performed with the Phy (https://github.com/cortex-lab/phy) software.

All analyses were performed using custom made scripts on Matlab (Mathworks). Statistical significance was considered at p<0.05.

## Acknowledgements

This work was supported by the French National Research Agency (ANR-22-CE17-0016) European Research Council under the European Union’s Horizon 2020 research and innovation program grant #646925 and #951330 (HOPE), the Fondation Bettencourt Schueller, and the Fondation Allianz-Institut de France. S.M. was funded by the “Ministère de l’Enseignement Supérieur, de la Recherche et de l’Innovation”. We would like to thank all INMED members for scientific discussions about the project; H Rouault, S. Sarno, R. Khazipov, M. Minlebaev, M. Picardo, P Quilichini, J Epsztein and J Koenig for their technical, computational and scientific assistance throughout the project. We are grateful to members of the Khazipov team for sharing equipment and facilities, A. Carabalona, A. Fortoul, and E. Buhler for their training in-utero injections; F. Michel from the INMED imaging facility (InMagic), M. Kurz for her assistance with the maintenance and generation of transgenic mice; A. Monthiel and F. Bader from the INMED cellular molecular biology platform for express genotyping, the staff of the INMED animal facility, G. Gay from the Turing Centre for living systems multi-engineering platform for custom software design and R. Martinez for his logistical support.

---

**Supplementary Figure 1.**
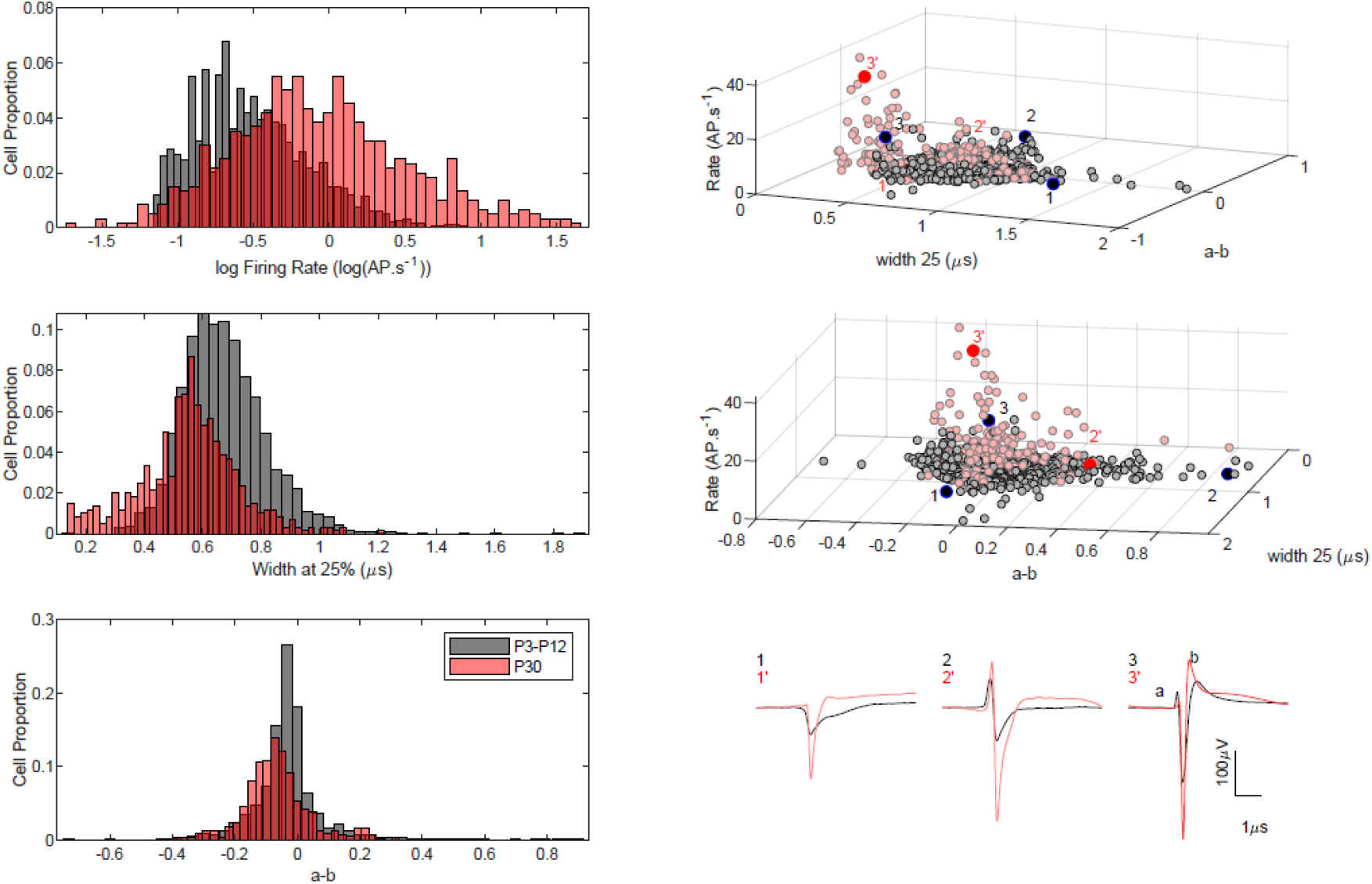
Left : histograms representing the proportion if single unit firing properties in immature (P3-P12, grey) vs. juvenile mice (P27-P30, red). Bottom. a and b refer to the pre and post hyperpolarization observed in single units as indicated in the right-side panel. Right: scatterplors of single unit properties recorded in both age groups. Juvenile fast spiking GABAergic cells form of a cluster in the high rate, low width corner. The cluster is absent in neonates. Bottom traces: average waveforms of cells indicated above in black (neonates) and red (juvenile) for waveforms with a large width (left), waveforms with a prominent biphasic spike (middle -see Someck et al 2023) and typical fast-spiking waveforms (right).

**Supplementary Figure 2:**
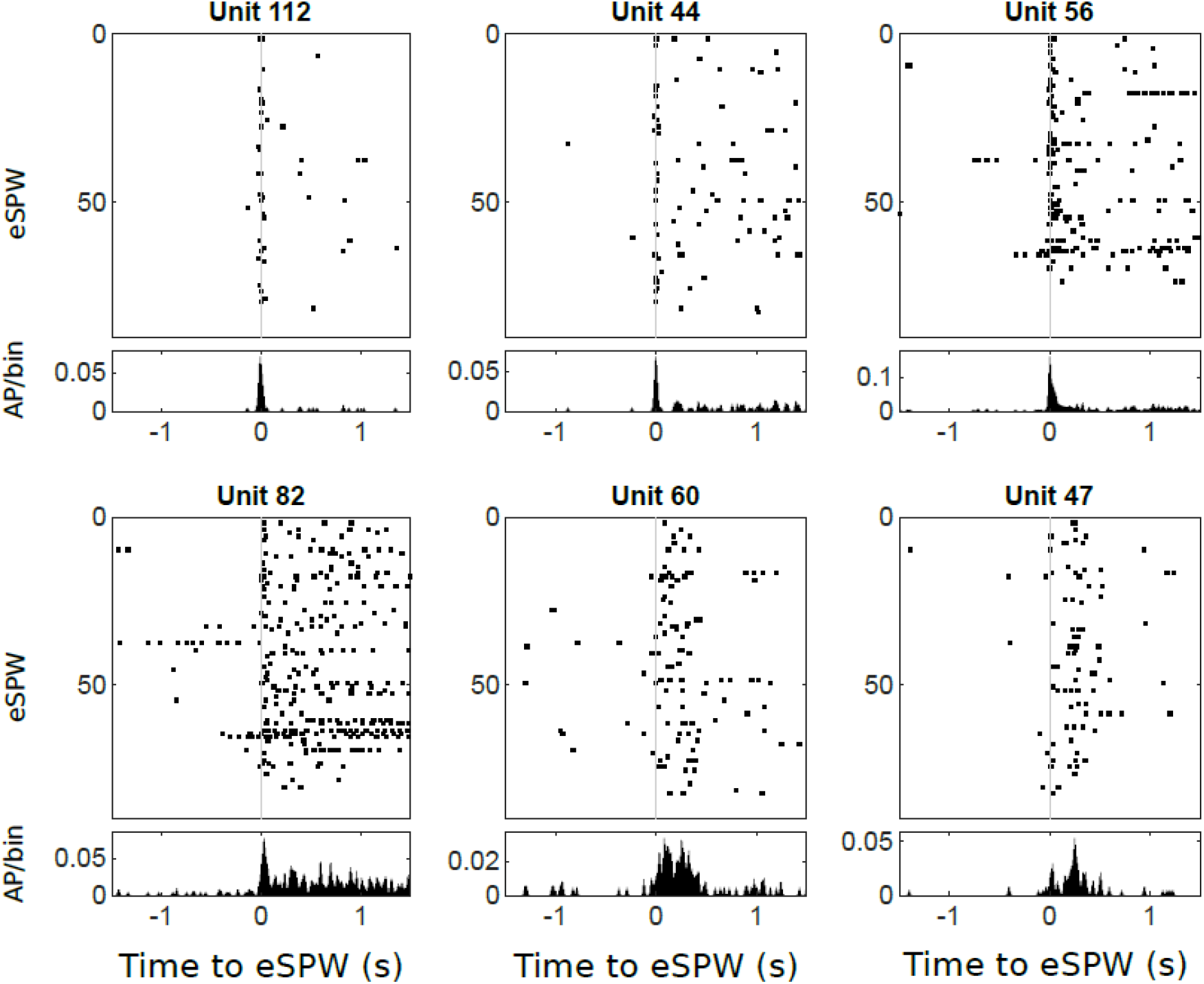
Rasters and corresponding histograms of 6 simultaneously recorded cells in a P4 mouse. While there is a variety of cell response types, each cell tend to fire in a stereotypical way following eSPWs.

**Supplementary Figure 3:**
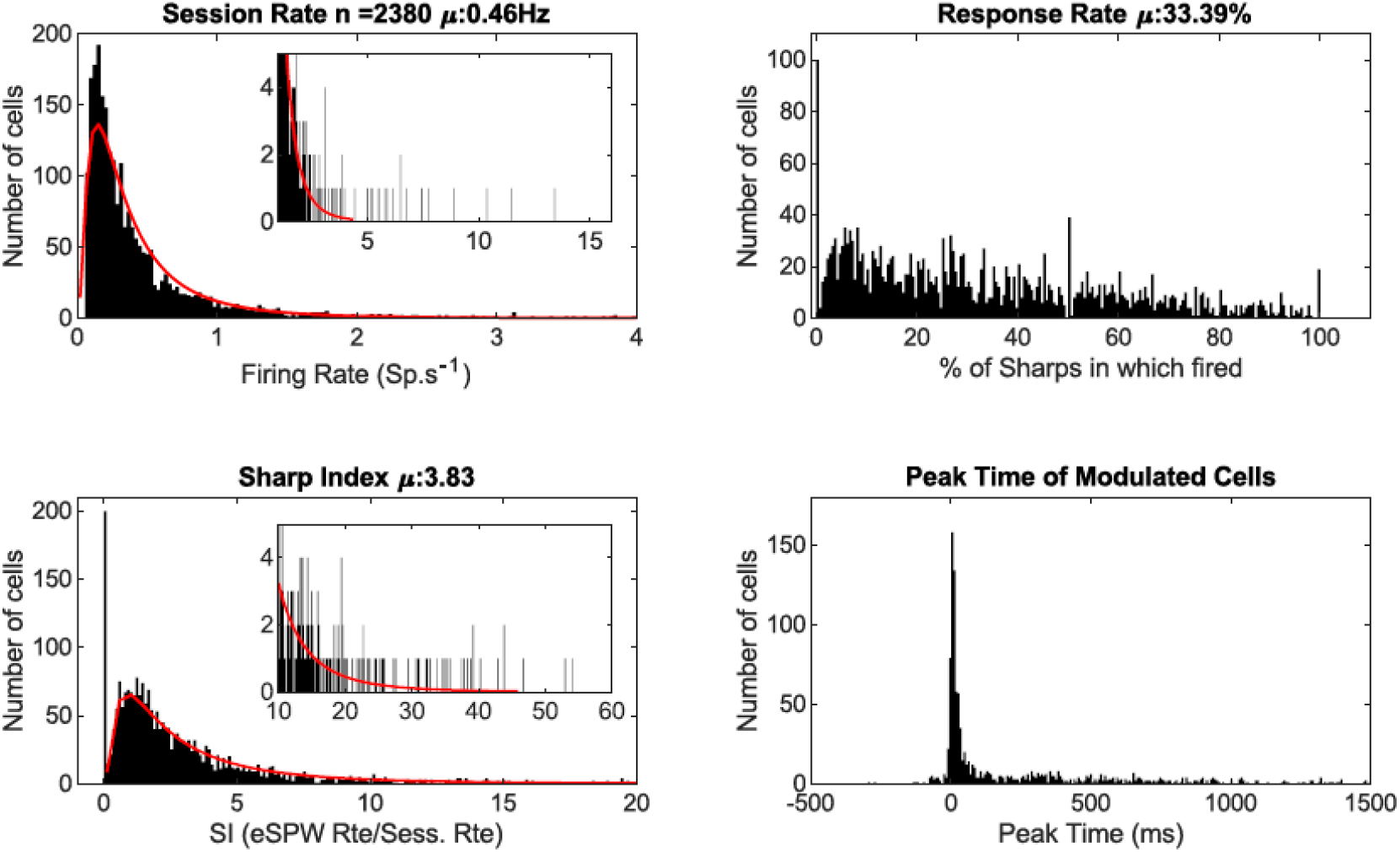
Histograms of cell discharge properties

**Supplementary Figure 4:**
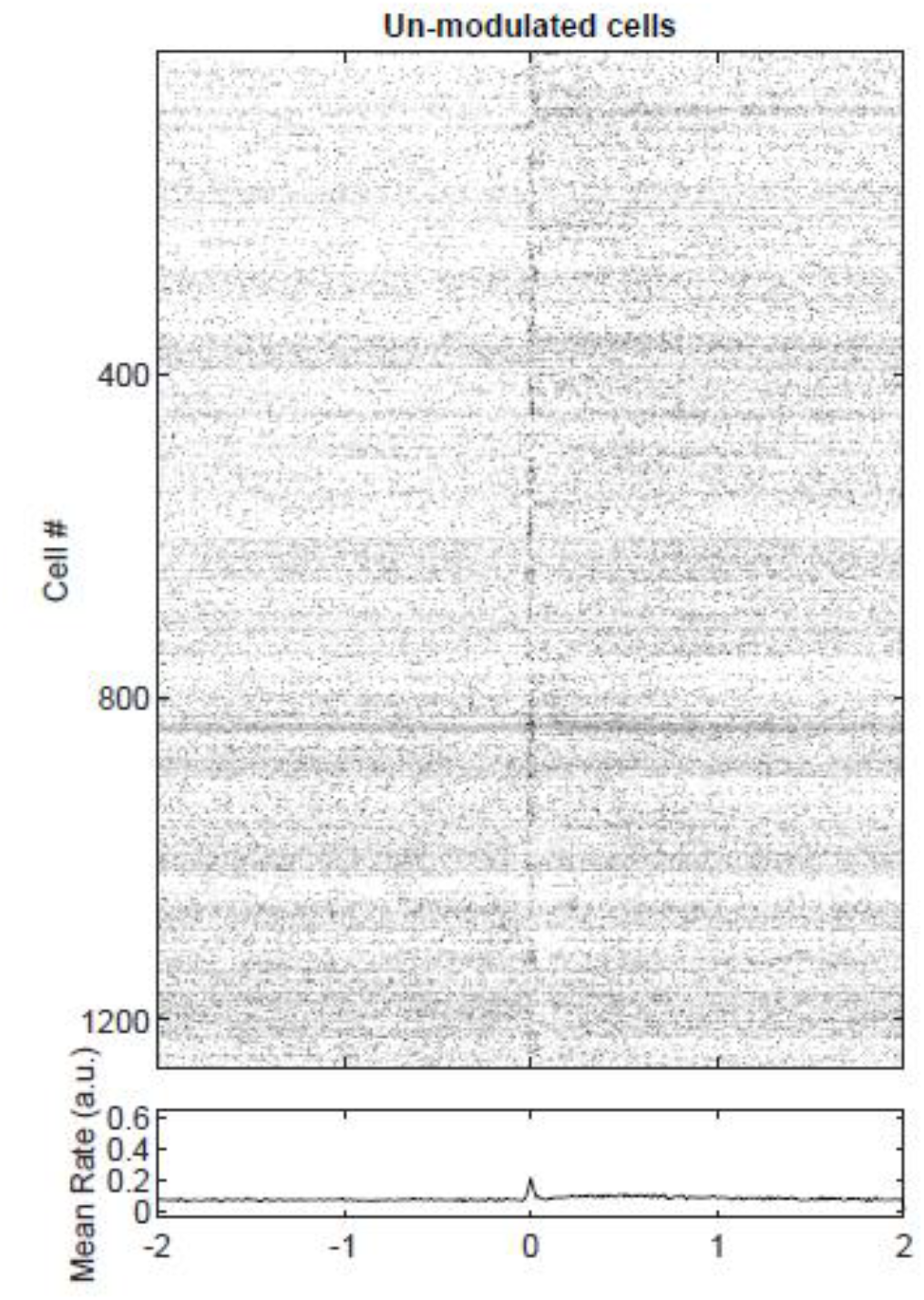
PETHs for un-modulated cells. Although cells did not reach significance of a systematic activation during eSPWs, an excess firing is visible for all cells at the time of eSPW trough.

**Supplementary Figure 5:**
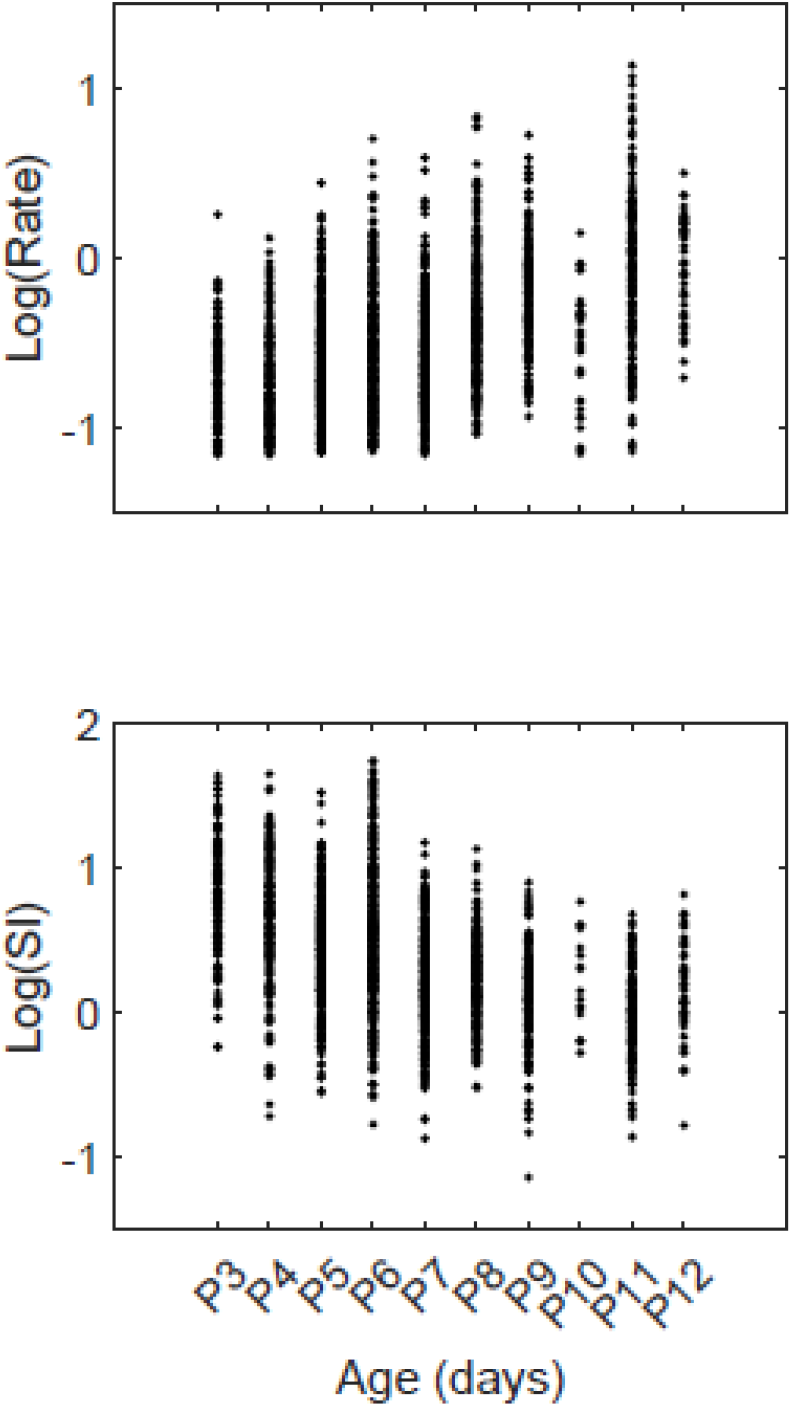
Relationship between age and session rate or SI (log values) for all recorded cells. Each dot corresponds to a cell.

**Supplementary Figure 6:**
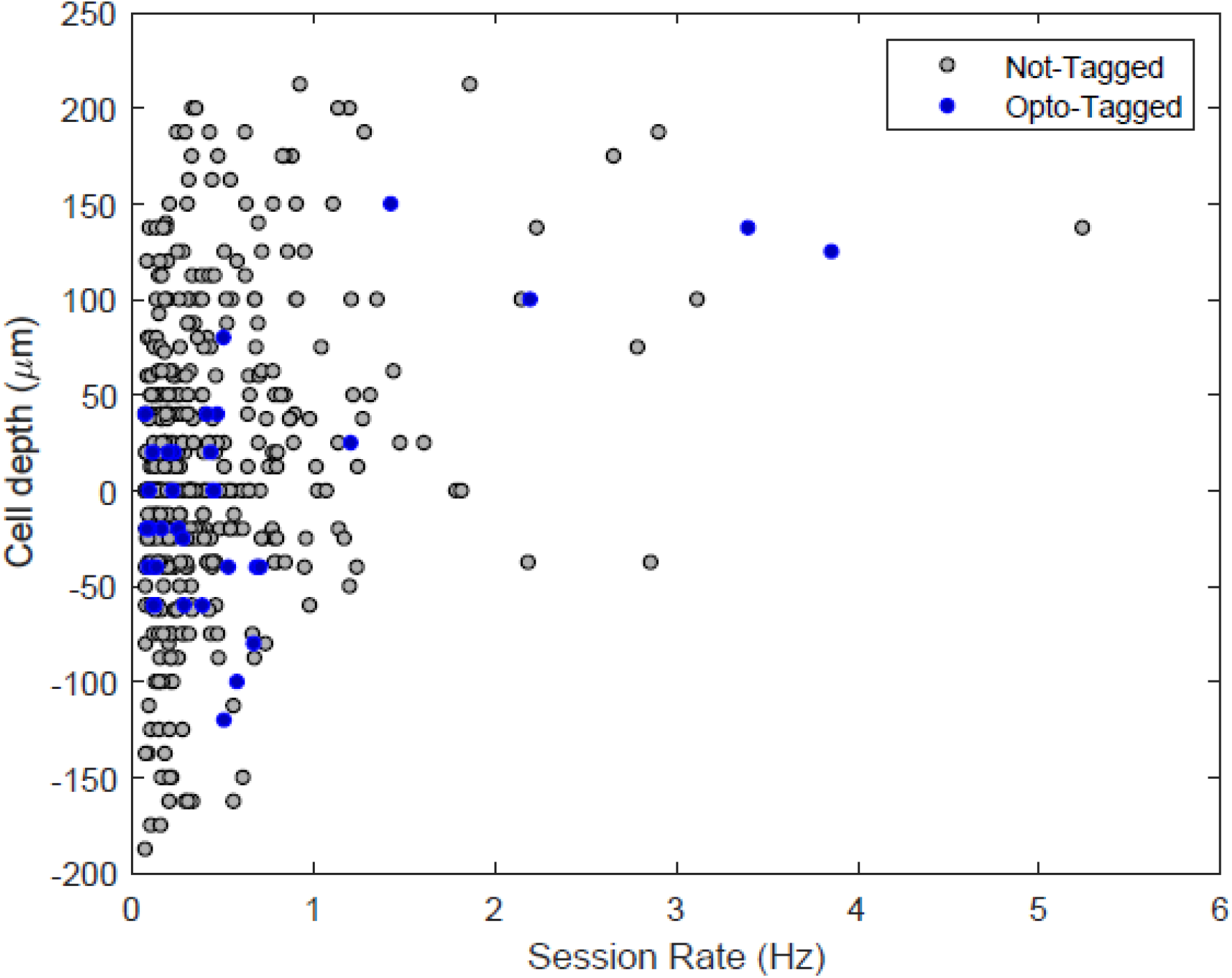
Session rate vs. localization in the radial axis of opto-tagged and not-tagged neurons. Each dot is a single neuron. Cell localization is estimated from histology, the presence of APs on electrode contacts and the site of inversion of eSPW LFP (zero-value, which is estimated to be on the superficial edge of the pyramidal cell layer).

