## Supplementary Figures for "GABAergic neurons are major contributors of network inhibition in the neonatal hippocampus in-vivo"

### Supplementary data

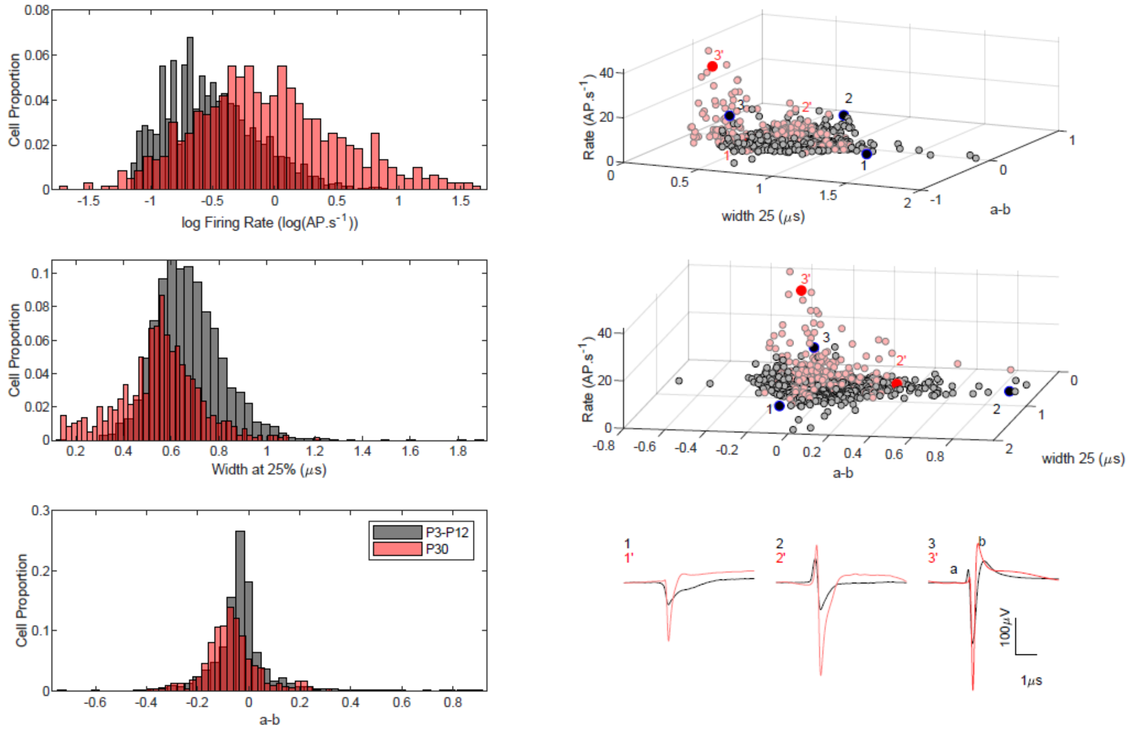


Supplementary Figure 1. Left : histograms representing the proportion if single unit firing properties in immature (P3-P12, grey) vs. juvenile mice (P27-P30, red). Bottom. a and b refer to the pre and post hyperpolarization observed in single units as indicated in the right-side panel. Right: scatterplors of single unit properties recorded in both age groups. Juvenile fast spiking GABAergic cells form of a cluster in the high rate, low width corner. The cluster is absent in neonates. Bottom traces: average waveforms of cells indicated above in black (neonates) and red (juvenile) for waveforms with a large width (left), waveforms with a prominent biphasic spike (middle -see Someck et al 2023) and typical fast-spiking waveforms (right).

reference: Someck, S., Levi, A., Sloin, H.E. *et al.* Positive and biphasic extracellular waveforms correspond to return currents and axonal spikes. *Commun Biol* **6**, 950 (2023). https://doi.org/10.1038/s42003-023-05328-6

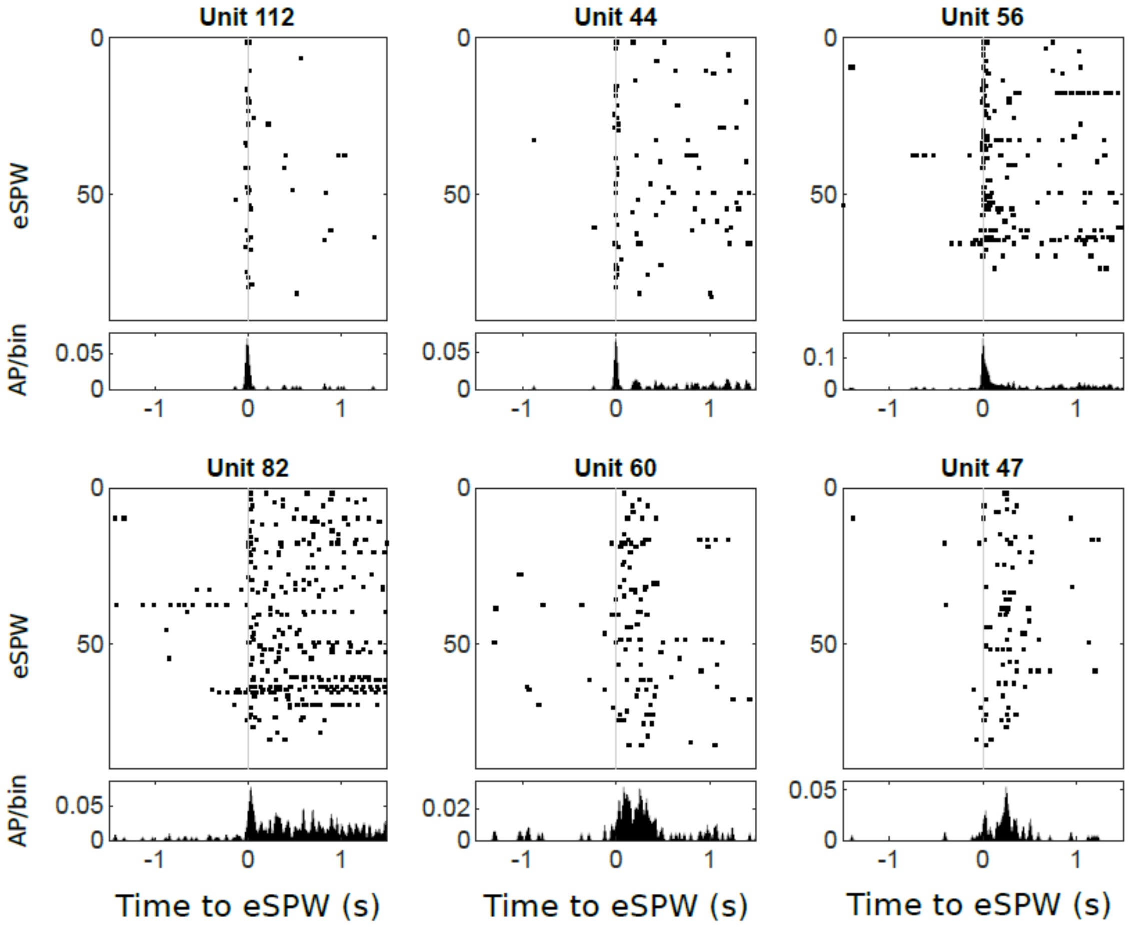


Supplementary Figure 2: Rasters and corresponding histograms of 6 simultaneously recorded cells in a P4 mouse. While there is a variety of cell response types, each cell tend to fire in a stereotypical way following eSPWs.

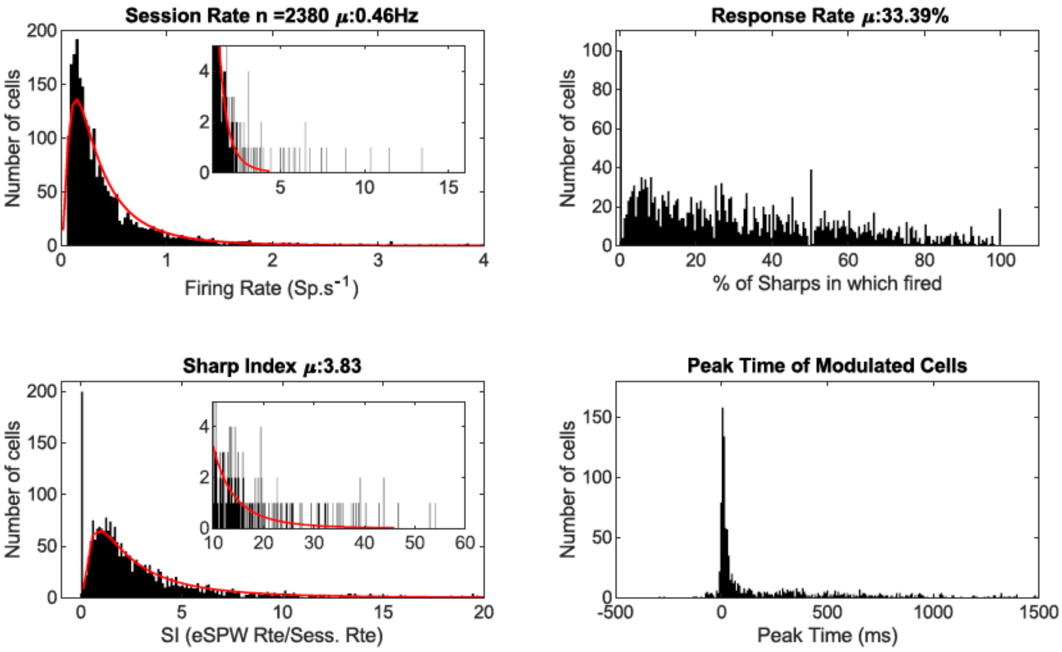


Supplementary Figure 3: Histograms of cell discharge properties

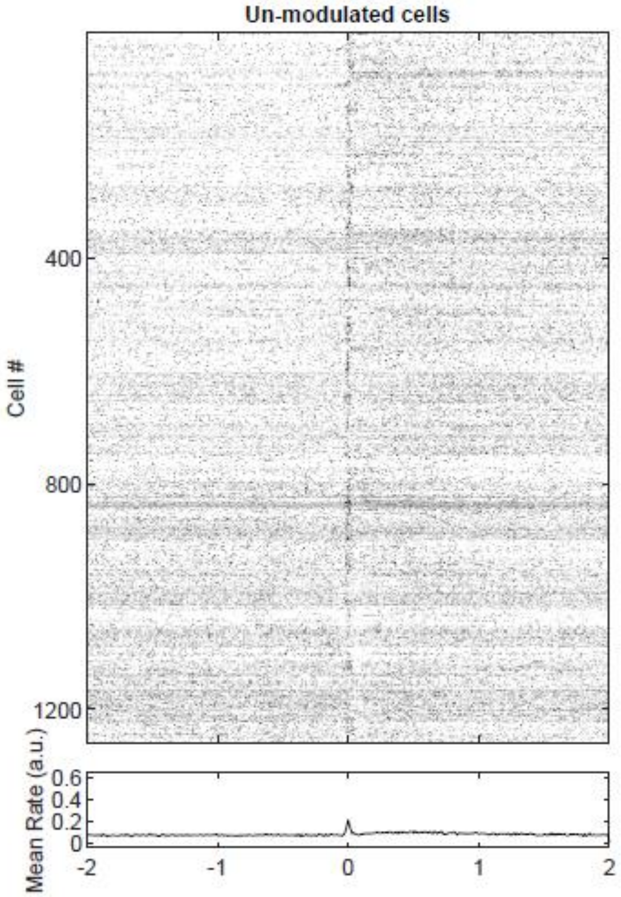


Supplementary Figure 4: PETHs for un-modulated cells. Although cells did not reach significance of a systematic activation during eSPWs, an excess firing is visible for all cells at the time of eSPW trough.

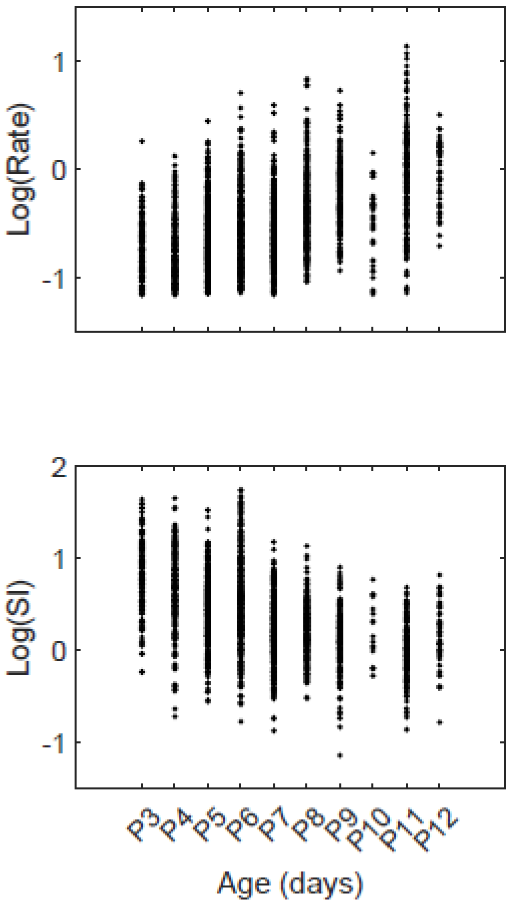


Supplementary Figure 5: Relationship between age and session rate or SI (log values) for all recorded cells. Each dot corresponds to a cell.

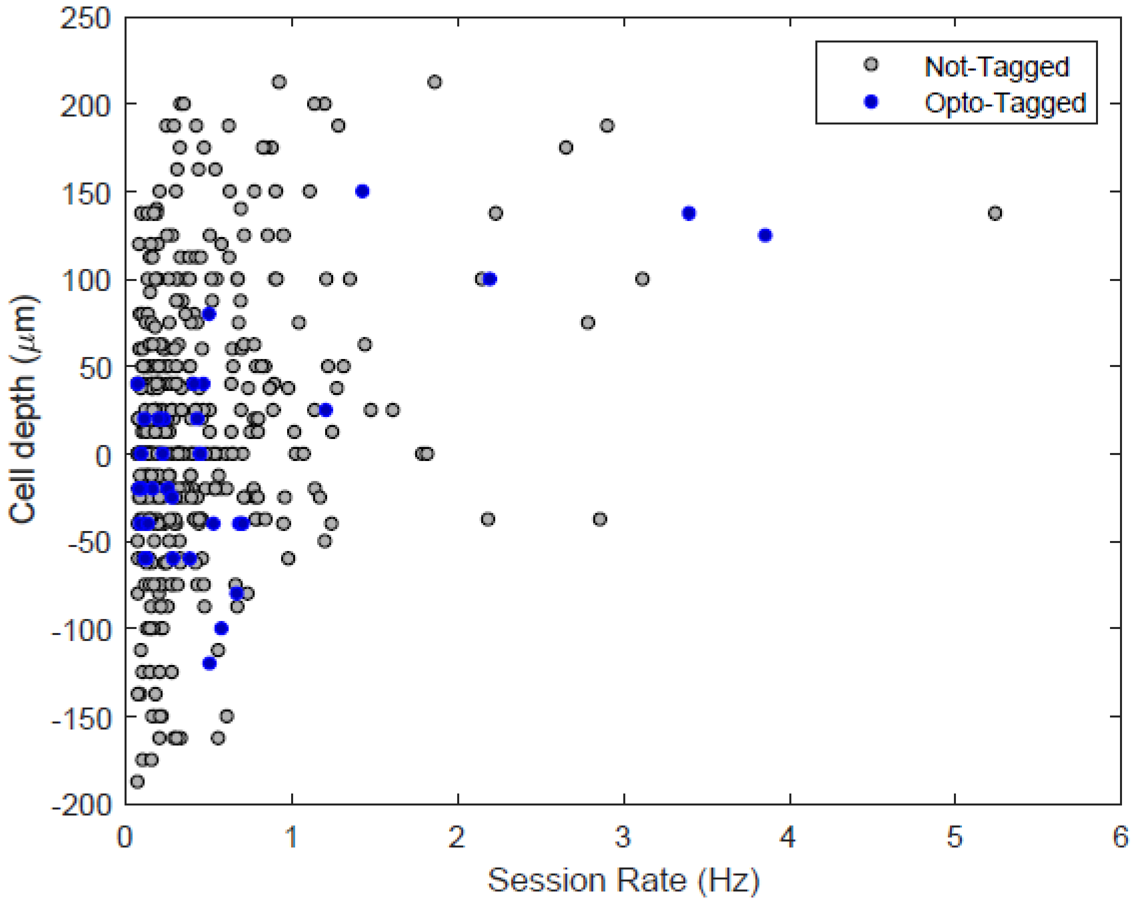


Supplementary Figure 6: Session rate vs. localization in the radial axis of opto-tagged and not-tagged neurons. Each dot is a single neuron. Cell localization is estimated from histology, the presence of APs on electrode contacts and the site of inversion of eSPW LFP (zero-value, which is estimated to be on the superficial edge of the pyramidal cell layer).
